# MicroScope: ChIP-seq and RNA-seq software analysis suite for gene expression heatmaps

**DOI:** 10.1101/034694

**Authors:** Bohdan B. Khomtchouk, James R. Hennessy, Claes Wahlestedt

## Abstract

We propose a user-friendly ChIP-seq and RNA-seq software suite for the interactive visualization and analysis of genomic data, including integrated features to support differential expression analysis, interactive heatmap production, principal component analysis, gene ontology analysis, and dynamic network analysis.

MicroScope is hosted online as an R Shiny web application based on the D3 JavaScript library: http://microscopebioinformatics.org/. The methods are implemented in R, and are available as part of the MicroScope project at: https://github.com/Bohdan-Khomtchouk/Microscope.

## Background

Most currently existing heatmap software produce static heatmaps (Saeed et al. 2003, Reich et al. 2006, Verhaak et al. 2006, Qlucore, GENE-E, Chu et al. 2008, Khomtchouk et al. 2014), without features that would allow the user to dynamically interact with, explore, and analyze the landscape of a heatmap via integrated tools supporting user-friendly analyses in differential expression, principal components, gene ontologies, and networks. Such features would allow the user to engage the heatmap data in a visual and analytical manner while in real-time, thereby allowing for a deeper, quicker, and more comprehensive data exploration experience.

An interactive, albeit non-reproducible heatmap tool was previously employed in the study of the transcriptome of the *Xenopus tropicalis* genome (Tan et al. 2013). Likewise, manual clustering of dot plots depicting RNA expression is an integral part of the Caleydo data exploration environment (Turkay et al., 2014). Chemoinformatic-driven clustering can also be toggled in the user interface of Molecular Property Explorer (Kibbey and Calvet, 2005). Furthermore, an interactive heatmap software suite was previously developed with a focus on cancer genomics analysis and data import from external bioinformatics resources (Perez-Llamas & Lopez-Bigas, 2011). Most recently, a general-purpose heatmap software providing support for transcriptomic, proteomic and metabolomic experiments was developed using the R Shiny framework (Babicki et al. 2016).

Moreover, an interactive cluster heatmap library, InCHlib, was previously proposed for cluster heatmap exploration (Škuta et al. 2014), but did not provide built-in support for gene ontology, principal component, or network analysis. However, InCHlib concentrates primarily in chemoinformatic and biochemical data clustering analysis, including the visualization of microarray and protein data. On the contrary, MicroScope is designed specifically for ChIP-seq and RNA-seq data visualization and analysis in the differential expression, principal component, gene ontology, and network analysis domains. In general, prior software has concentrated primarily in hierarchical clustering, searching gene texts for substrings, and serial analysis of genomic data, with no integrated features to support the aforementioned built-in features (Saldanha 2004, Caraux and Pinloche 2005, Wu et al. 2010).

As of yet, no free, open-source heatmap software has been proposed to explore heatmaps at such multiple levels of genomic analysis and interactive visualization capacity. Here we propose a user-friendly genome software suite designed to handle dynamic, on-the-fly JavaScript visualizations of gene expression heatmaps as well as their respective differential expression analysis, principal component analysis, gene ontology analysis, and network analysis of genes.

## Implementation

MicroScope is hosted online as an R Shiny web server application. MicroScope may also be run locally from within R Studio, as shown here: https://github.com/Bohdan-Khomtchouk/Microscope. MicroScope leverages the cumulative utility of R's d3heatmap (Cheng et al. 2015), shiny (Chang et al. 2015), stats (R Core Team, 2015), htmlwidgets (Vaidyanathan et al. 2015), RColorBrewer (Neuwirth, 2014), dplyr (Wickham et al. 2015), data.table (Dowle et al. 2015), goseq (Young et al. 2010), GO.db (Carlson, 2016a), and networkD3 (Gandrud et al. 2015) libraries to create an integrative web browser-based software experience requiring absolutely no programming or statistical experience from the user, or even the need to download R on a local computer.

MicroScope employs the Bioconductor package edgeR (Robinson et al. 2010) to create a one-click, built-in, user-friendly differential expression analysis feature that provides differential expression analysis of gene expression data based on the quantile-adjusted conditional maximum likelihood (qCML) procedure and the Benjamini & Hochberg correction. edgeR is a count-based statistical method that expects input data in the form of a matrix of integer values. The value in the *i*-th row and the *j*-th column of the matrix tells how many reads (or fragments, for paired-end RNA-seq) have been unambiguously assigned to gene *i* in sample *j* (Love et al. 2016). Analogously, for other types of assays, the rows of the matrix might correspond e.g., to binding regions (with ChIP-seq), species of bacteria (with metagenomic datasets), or peptide sequences (with quantitative mass spectrometry). In general, the values in the matrix must be raw counts of sequencing reads/fragments. This is important for the statistical model to hold, as only the raw counts allow assessing the measurement precision correctly. It is important to never provide counts that were pre-normalized for sequencing depth/library size, as the statistical model is most powerful when applied to raw counts, and is designed to account for library size differences internally via a series of built-in normalization procedures.

The edgeR results supply the user with rank-based information about nominal p-value, false discovery rate, fold change, and counts per million in order to establish which specific genes in the data are differentially expressed with a high degree of statistical significance. This information, in turn, is used to investigate the top gene ontology categories of differentially expressed genes, which can then be conveniently visualized as interactive network graphics. Finally, MicroScope provides user-friendly support for principal component analysis via the generation of biplots, screeplots, and summary tables. PCA is supported for both covariance and correlation matrices via R’s prcomp() function in the stats package.

## Results & Discussion

Figure 1 shows the MicroScope user interface (UI) upon login. After a user inputs an RNA-seq/ChIP-seq data file containing read counts per gene per sample, the user is guided through the differential expression analysis (Figure 2) which, in turn, leads to the heatmap visualization stage of differentially expressed genes at user-specified statistical cutoff parameters (Figure 3). Heatmaps visualizing statistically significant genes, as determined by the differential expression analysis, can be customized in a variety of ways, through user-friendly methods such as:

- Statistical parameters visualization cutoff widget (p-value and/or FDR)
- *log*_2_ data transformation widget
- Multiple heatmap color schemes widget
- Hierarchical clustering widget
- Row/column dendrogram branch coloring widget
- Row/column font size widget
- Heatmap download widget

**Figure 1.**
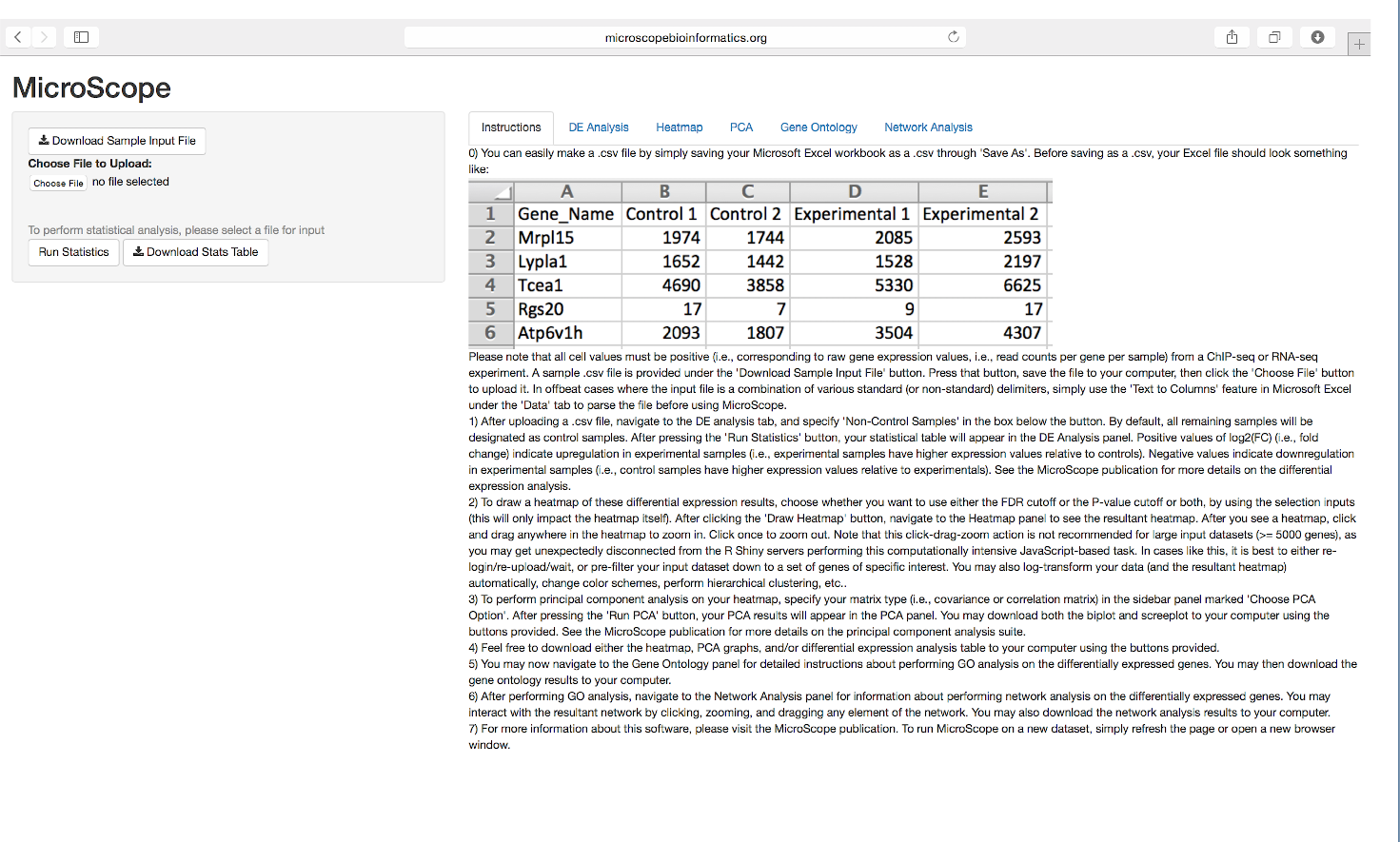
MicroScope user interface. MicroScope UI shown at login, showcasing the Instructions tab and differential expression analysis feature, as well as features such as: sample file download, input file upload, ‘Run Statistics’ widget, and ‘Download Stats Table’ widget. Additional UI features are sequentially unlocked as the user progresses through the MicroScope software suite.

**Figure 2.**
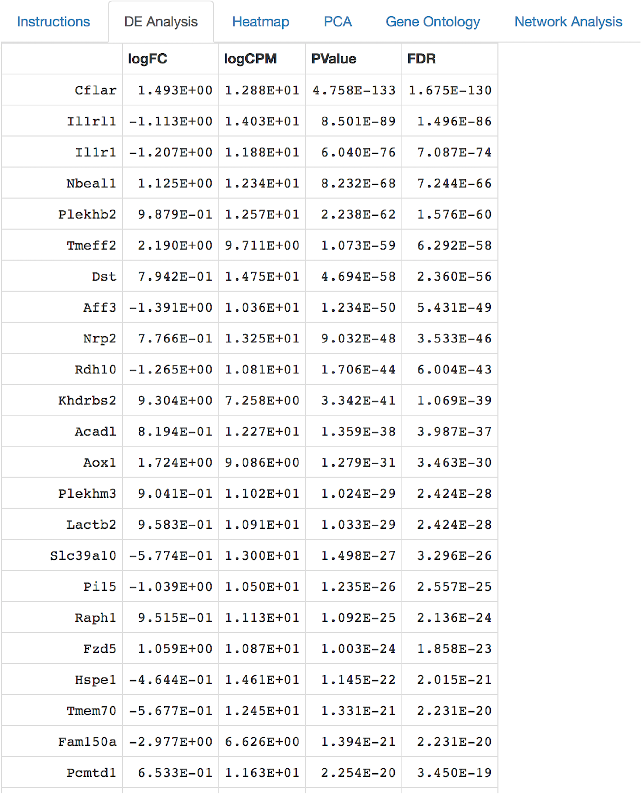
Differential expression analysis tabulated results. Once the input data is uploaded, a quantile-adjusted conditional maximum likelihood (qCML) procedure and the Benjamini-Hochberg correction are used to supply the user with information about the nominal p-value, false discovery rate, fold change, and counts per million calculations for differentially expressed genes. The edgeR package is used.

**Figure 3.**
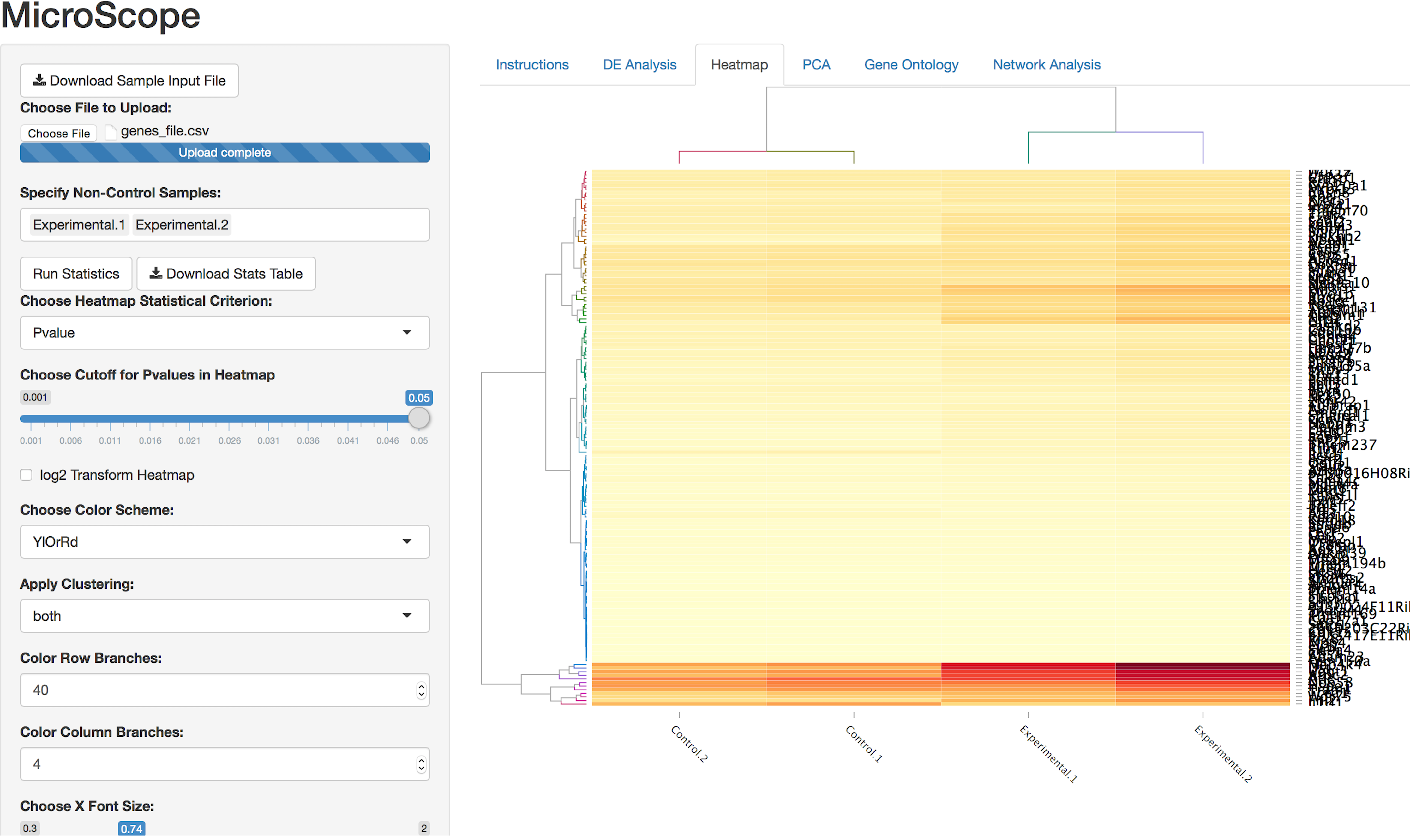
Interactive heatmap visualization. MicroScope heatmap options showcasing the magnification feature as well as features such as: statistical parameter settings, *log*_2_ data transformation, multiple heatmap color schemes, hierarchical clustering, row/column dendrogram branch coloring, row/column font size, and heatmap download button.

MicroScope allows the user to magnify any portion of a heatmap by a simple click-and-drag feature to zoom in, and a click-once feature to zoom out. MicroScope is designed with large gene expression heatmaps in mind, where individual gene labels overlap and render the text unreadable. However, MicroScope allows the user to repeatedly zoom in to any sector of the heatmap to investigate a region, cluster, or even a single gene. MicroScope also allows the user to hover the mouse pointer over any specific gene to show gene name, expression level, and column ID. It should be noted that specifying the heatmap statistical parameters impacts the contents of the heatmap visualization itself, as stringent cutoffs will naturally result in less genes displayed. However, the downstream PCA or gene ontology or network analysis is not impacted by these heatmap visualizations. In other words, all downstream analyses are performed on the entire input dataset. It should also be noted that prior to visualizing heatmaps in MicroScope, experiment-specific data normalization procedures are left to the discretion of the user (Conesa et al. 2016, Soneson & Delorenzi 2013, Bailey et al. 2013, Shin et al. 2013), depending on whether the user wants to visualize differences in magnitude among genes or see differences among samples.

One of the user-friendly features within MicroScope is that it is responsive to the demands asked of it by the user. For example, gene ontology analysis buttons are not provided in the UI until a user runs differential expression analysis, which constitutes a prerequisite step required prior to conducting a successful gene ontology analysis. In other words, MicroScope is user-responsive in the sense that it automatically unlocks new features only as they become needed when the user progresses through successive stages in the software. Furthermore, MicroScope automatically provides short and convenient written guidelines directly in the UI to guide the user to the next steps in the usage of the software. As such, complex analytical operations can be performed by the user in a friendly, step-by-step fashion, each time facilitated by the help of the MicroScope software suite, which adjusts to the needs of the user and provides written guidelines on the next steps to pursue. It should be noted that the differential expression analysis in MicroScope (qCML and Benjamini & Hochberg correction) is broadly applicable to be run on any ChIP-seq or RNA-seq data inputted by the user.

Following the successful completion of the differential expression analysis and interactive heatmap visualization, a user is automatically supplied a suite of UI widgets to perform principal component analysis. The user is given the choice to specify the matrix type (i.e., covariance or correlation matrix) in the sidebar panel marked’Choose PCA Option’. After the PCA is completed, the user is supplied with a biplot and screeplot to visualize the results, as well as tabulated information showing the relative importance of each principal component.

Following the successful completion of the PCA (Figure 4), the user is prompted with more UI widgets to proceed to the gene ontology analysis. Specifying values for these features and clicking the Do Gene Ontology Analysis button returns a list of the top gene ontology (GO) categories according to these exact specifications set by the user (Figure 5). Supported organisms for GO category analysis include: human (Carlson, 2016b), mouse (Carlson, 2016c), rat (Carlson, 2016d), zebrafish (Carlson, 2016e), worm (Carlson, 2016f), chimpanzee (Carlson, 2016g), fly (Carlson, 2016h), yeast (Carlson, 2016i), bovine (Carlson, 2016j), canine (Carlson, 2016k), mosquito (Carlson, 2016l), rhesus monkey (Carlson, 2016m), frog (Carlson, 2016n), and chicken (Carlson, 2016o).

**Figure 4.**
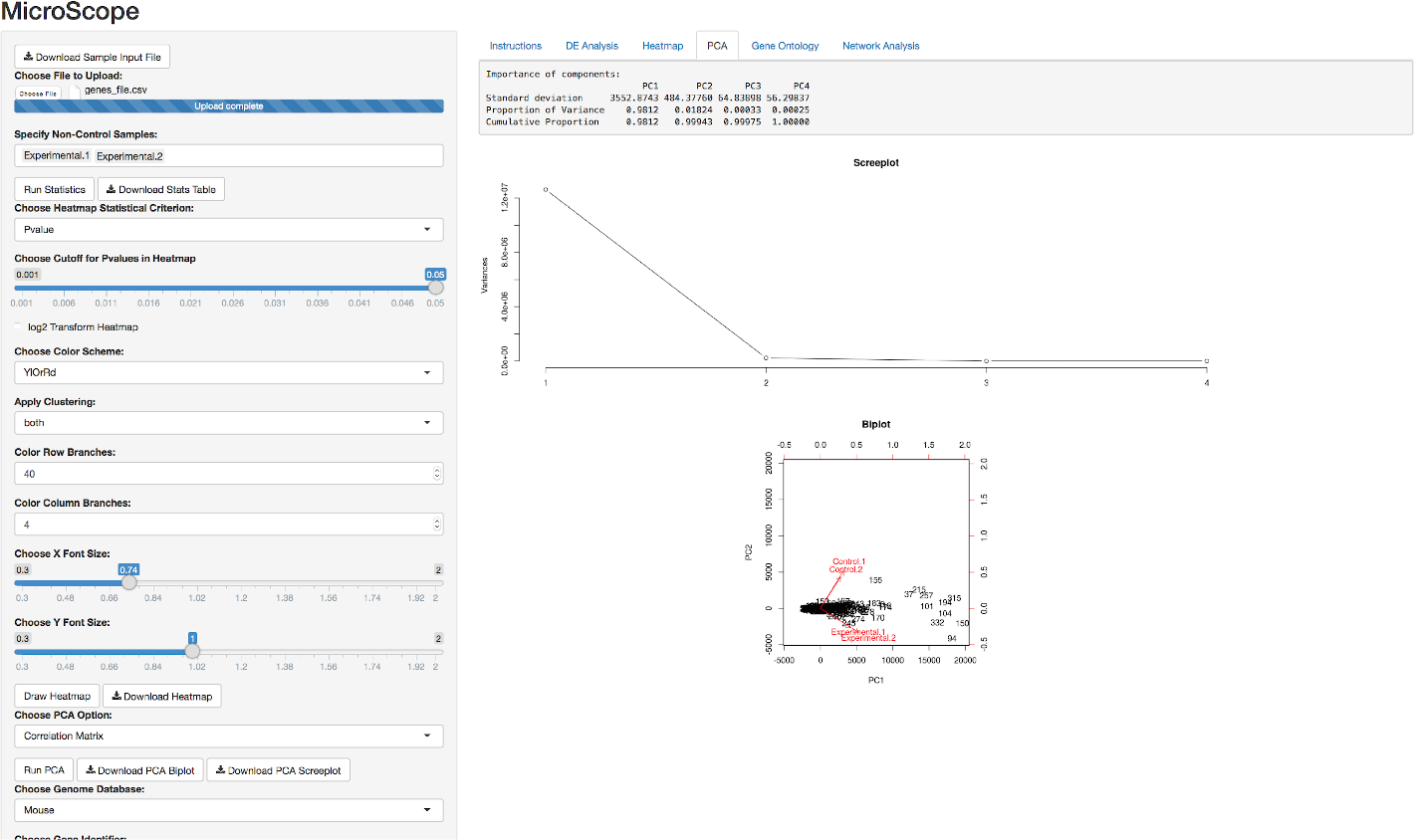
Principal component analysis. Tabulated summary table of importance of principal components, as well as biplot and screeplot graphics visualizations, are produced to investigate variation and patterns in gene expression.

**Figure 5.**
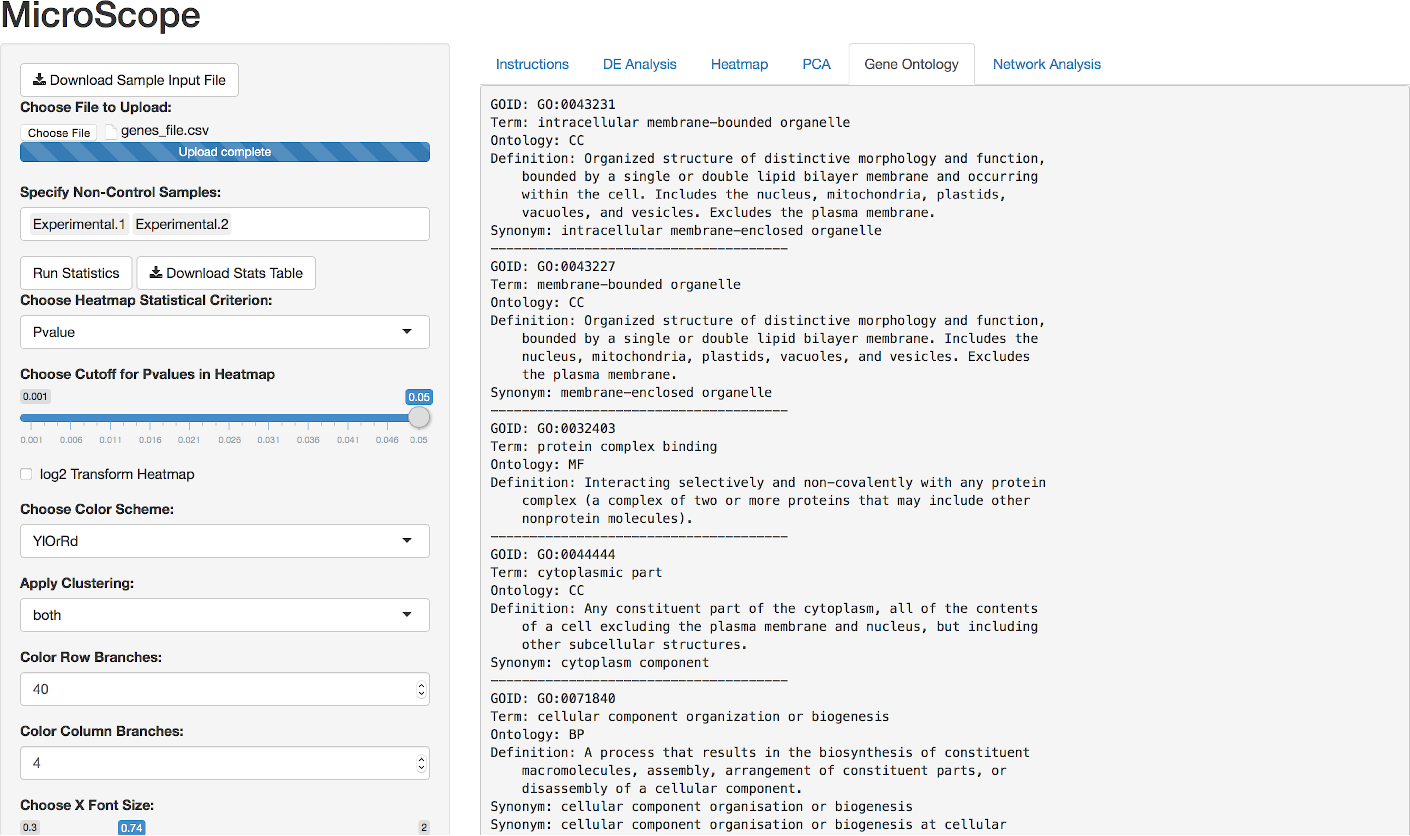
Gene ontology analysis tabulated results. Top gene ontology categories are automatically calculated and returned as a ranked list in the UI.

The successful completion of this step can be followed up by running a network analysis on the top GO categories, thereby generating network graphics corresponding to the number of top gene ontology categories previously requested by the user (Figure 6). Nodes represent either gene names or gene ontology identifiers, and links represent direct associations between the two entities. In addition to serving as a visualization tool, this network analysis capability automatically identifies differentially expressed genes that are present within each top gene ontology, which is a level of detail not readily available by running gene ontology analysis alone. By immediately extracting the respective gene names from each top gene ontology category, MicroScope’s network analysis features serve to aid the biologist in identifying the top differentially expressed genes in the top respective gene ontology categories. Figure 7 compares interactive network visualizations of the top two gene ontologies, thereby demonstrating the immediate responsiveness of MicroScope’s network graphics to user-specified settings (e.g., number of top gene ontologies to display widget).

**Figure 6.**
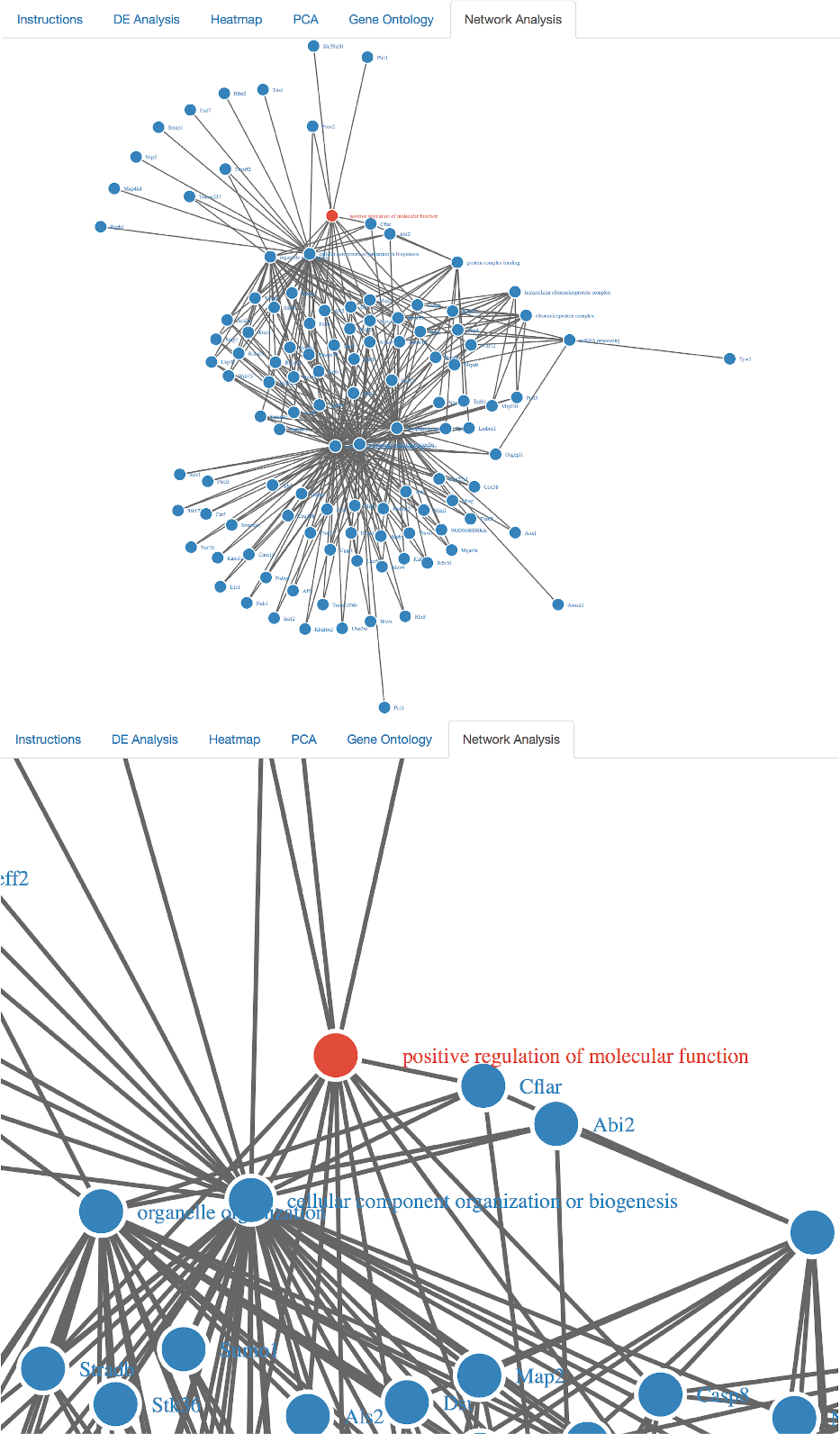
Network graphics visualizations of top gene ontology categories. Differentially expressed genes belonging to the respective gene ontology categories are automatically displayed during the network analysis of the data. Top ten gene ontologies (and their respective genes) are shown here. Networks are zoomable and dynamically interactive, allowing the user to manually drag nodes across the screen to explore gene_name-gene_ontology interconnectivity and network architecture.

**Figure 7.**
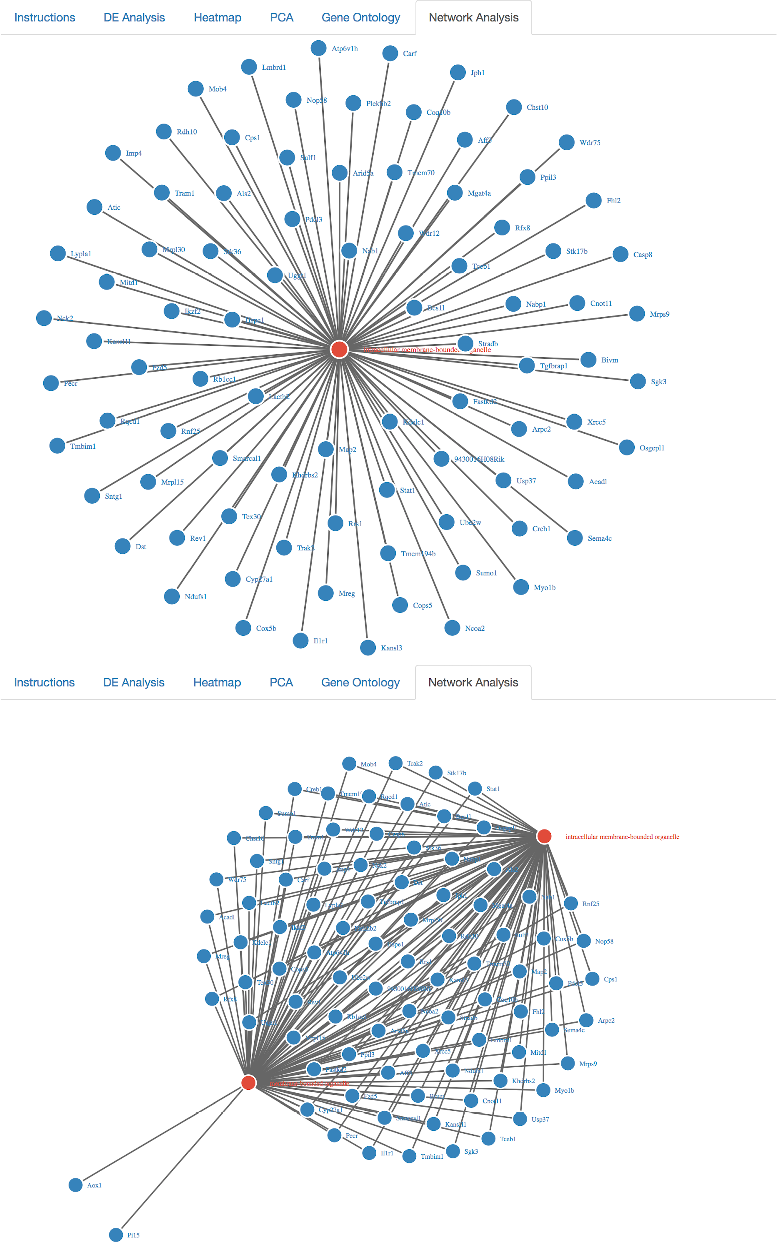
Network visualizations of first ranked gene ontology vs. top-two ranked gene ontologies. Comparison of dynamically interactive network graphics at various user-specified gene ontology settings (e.g., ‘Choose How Many Top Gene Ontologies to Display’ button in the UI). Clearly, the GO category “membrane-bounded organelle” contains two unique genes, while the rest are (perhaps unsurprisingly) shared in common with the GO category “intracellular membrane-bounded organelle”.

## Conclusion

We provide access to a user-friendly web application designed to visualize and analyze dynamically interactive heatmaps within the R programming environment, without any prerequisite programming skills required of the user. Our software tool aims to enrich the genomic data exploration experience by providing a variety of complex visualization and analysis features to investigate gene expression datasets. Coupled with a built-in analytics platform to pinpoint statistically significant differentially expressed genes, an interactive heatmap production platform to visualize them, a principal component analysis platform to investigate variation and patterns in gene expression, a gene ontology platform to categorize the top gene ontology categories, and a network analysis platform to dynamically visualize gene ontology categories at the gene-specific level, MicroScope presents a significant advance in heatmap technology over currently available software.

## Author’s contributions

BBK conceived of the study. BBK, JRH, and VDR wrote the code. CW participated in the management of the source code and its coordination. BBK wrote the paper. All authors read and approved the final manuscript.

## Acknowledgements

BBK dedicates this work to the memory of his uncle, Taras Khomchuk. BBK wishes to acknowledge the financial support of the United States Department of Defense (DoD) through the National Defense Science and Engineering Graduate Fellowship (NDSEG) Program: this research was conducted with Government support under and awarded by DoD, Army Research Office (ARO), National Defense Science and Engineering Graduate (NDSEG) Fellowship, 32 CFR 168a. CW thanks Vytas Dargis-Robinson for assistance in early stages of the project.

## Ethics

This study does not involve humans, human data or animals.

## Abbreviations used

FDR: false discovery rate
GO: gene ontology
UI: user interface
PCA: principal component analysis
DE: differential expression
qCML: quantile-adjusted conditional maximum likelihood

## Availability of Data and Materials

All source code has been made publicly available on Github at: https://github.com/Bohdan-Khomtchouk/Microscope.

## Figures as additional files

Figures have been uploaded as additional files. Standard BioMed Central bmc article LaTeX template has been used for production of figure captions.

